# Acceleration of Large-Scale Single Cell Differential Gene Expression Analysis with FastDE

**DOI:** 10.1101/2023.06.15.545003

**Authors:** Erin Connolly, Maneesha Aluru, Sriram Chockalingam, Vishal Dhere, Greg Gibson, Tony Pan

## Abstract

The major bottleneck in single cell transcriptomic processing is identification of marker genes for individual cell clusters. We introduce FastDE, an implementation of the standard Wilcoxon Rank Sum test in C++ that results in 50X speed-up of the standard Seurat pipeline on a dataset of 1.2 million cells. Proof of methodology is demonstrated by replicating age and gender effects on immune cell differentiation in two large PBMC atlases.

---

A critical step in single cell genomic profiling is to cluster cells and then identify marker genes that differentiate the clusters^1,2^. Marker identification is typically based on significance testing of the expression rank of transcript abundance among clusters^3^. Once identified, the markers can be used to annotate the identity or state of cell clusters, also providing a measure of robustness and repeatability across datasets. Expanded gene lists are often the basis for biological contrasts among treatments or effects of interest. Here we introduce an efficient algorithm for marker detection, FastDE, and demonstrate its use on a dataset of 2.7 million cells to ask how differentiation of peripheral blood immune cell-types changes with sex and age in humans.

One widely used open-source software for single cell transcriptome profiling is Seurat^4,5^. The typical workflow is to read in data, scale and transform, perform principal and canonical correlation analysis to identify cell clusters, project these onto 2-dimensional space with UMAP, and then compute Fold Change and Wilcoxon Rank Sum Tests to find cluster-specific marker genes. Seurat defaults to LIMMA^6^ for small datasets, but uses native R code for larger ones, and converts sparse to dense matrices, which places constraints on computational time and the size of dataset that can be analyzed.

Incremental versions of the software have improved clustering performance and incorporated options for differential expression analysis, but the FindMarkers function remains the rate limiting step for analysis.

We ran Seurat on a single core of a Xeon E7-8870 processor with publicly available datasets containing between 3,000 and 68,000 immune cells of varying complexity, and consistently observed that between 79% and 97% of the run-time was consumed by the FindMarkers function (**Supplementary Table 1**). The R language^7^ is limited to 2 billion non-zero elements in a sparse matrix and is therefore constrained for analyzing datasets with high cell counts. In addition, intermittent memory allocation failures were observed when using multiple cores with default Seurat parameters. To address these limitations, we recoded the key components of Seurat’s FindMarkers implementation in C++^8^ utilizing sparse matrix specific algorithms, large sparse matrix indices, and OpenMP^9^ to accelerate the workflow, thereby facilitating analysis of very large compendium datasets and improving multicore processing speed and stability.

Our FastDE software utilizes the Wilcoxon-Mann-Whitney U (Wilcoxon Rank Sum) test in C++ and is schematized in **Fig. 1**. For each gene, the observed counts are sorted once and the sorted rank is reused for comparing each cluster against all other clusters, similar to the approach of BioQC^10^. This is in contrast to Seurat, where the differentially expressed genes are identified by computing the full gene expression count matrix for each cluster, thus resulting in the same sort operation being repeated redundantly for every cluster. Applying this approach to both Seurat’s FoldChange and the Wilcoxon Rank Sum test, runtime, excluding file access, on the 68,000-cell dataset was reduced from 50 minutes to 6.5 minutes, nearly an 8-fold acceleration.

**Fig. 1:**
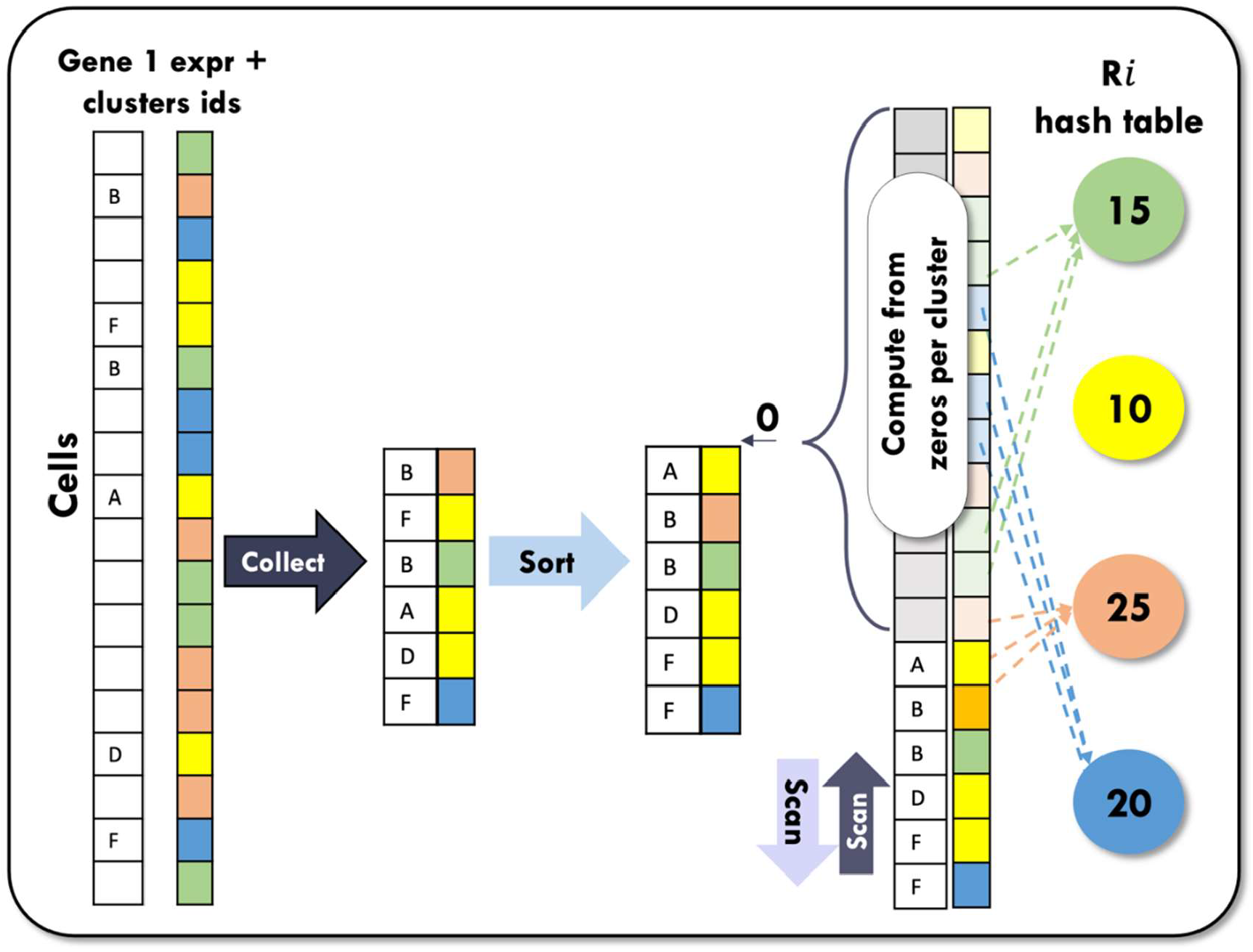
FastDE sparse matrix Workflow for implementing Wilcoxon Rank Sum test.

We further introduced three changes that collectively result in up to a 200-fold speed-up over Seurat for the FindMarkers step, and a 10-to 30-fold speed-up of the entire Seurat workflow, of which just 3% to 15% was taken up by the FastDE FindMarkers routine. First, we improved memory and cache usage through sequential access of intermediate arrays thus allowing hardware memory prefetching and use of a small cache-friendly hash table for accumulating rank sums. Second, ties correction factors were computed during rank summation via a linear scan of the array. Using the hash table and the tie correction factors, Rank Sum, and U-score statistics, the corresponding p-values are generated in constant time per cluster. These memory optimization steps halve the compute time for fold change and differential expression computation. Third, we leverage the sparsity of the count matrix directly in our FoldChange and Wilcoxon Rank Sum test algorithms. Whereas Seurat densifies a sparse matrix and processes both zero and non-zero entries, we leveraged the fact that all zero entries share the same rank. Their rank sum and tie correction factor contributions can be computed in constant time from the ranks of the non-zero values closest to zero and their counts. We confirmed that the FindMarkers output was consistent with Seurat on the same datasets. Elimination of the dense matrix, which typically contains 90% to 97% zero counts in droplet-based single cell RNAseq datasets, greatly expands the size of dataset that can be handled.

To confirm the utility of FastDE for standard differential expression analysis of very large datasets, we assembled a compendium of ∽2.7 million PBMCs from 678 adults sampled in 35 studies (Methods; **Fig. 1**). Clusters were identified in subsets of the data, and imported into a common Seurat object, focusing on nineteen common cell types robustly identified in all contributing datasets: naïve, intermediate and memory B cells, plasma cells, conventional and plasmocytoid dendritic cells, CD14 and CD16 monocytes, Cytotoxic CD4 T cells, NK cells, CD56+ NK cells, MAIT cells, central and effector memory and naive CD4+ and CD8+ T cells, and regulatory T cells (Tregs) **(Fig. 2d)**. We first performed FastDE on the entire dataset, and then partitioned the dataset into male and female subsets, each with age deciles, to investigate whether immune cells undergo age-related loss of cellular identity in a sex-specific manner.

**Figure 2:**
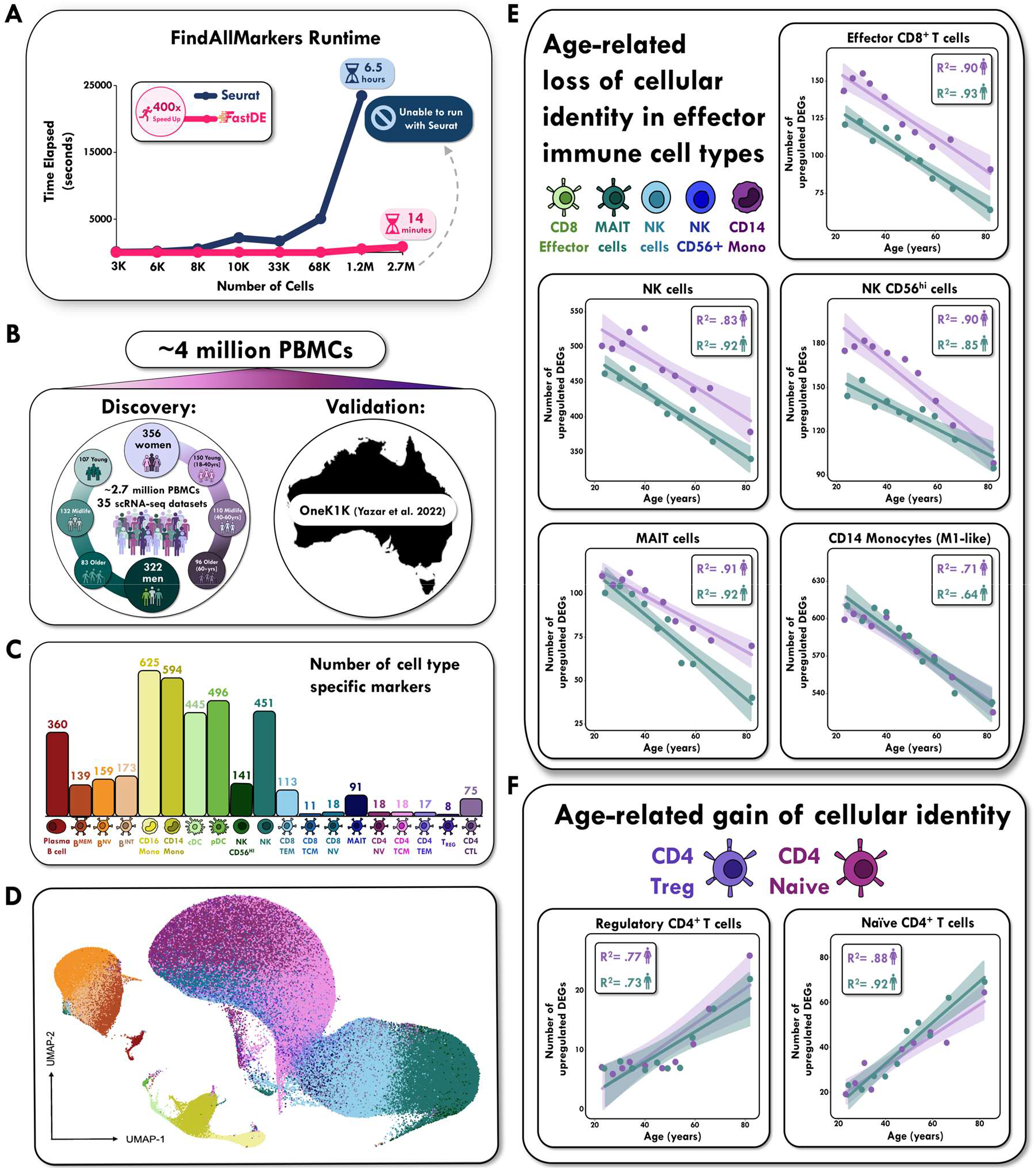
Study design and Summary of PBMC Atlas Results. **A**, Runtime benchmarks using Seurat’s base R wilcox.test (blue) and FastDE (pink). **B**, scRNA-seq data of PBMCs from a discovery cohort comprising of 356 women and 322 men (18-110 years old) and the independent 1.2 million-cell OneK1K validation atlas from 982 Australian adults (18-97 years old). **C**, Barplot showing the number of marker genes for each of 19 cell types considering all 2.74M cells in the discovery atlas. **D**, UMAP of PBMCs across all individuals in the discovery atlas, with 19 transcriptionally distinct populations. **E**, Positive correlation between age and the number of cell-type specific markers in effector immune cells (CD8 effector T cells; MAIT cells; NK cells; NK CD56+ cells; CD14 Monocytes). **F**, Negative correlation between age and the number of cell type specific markers in CD4 regulatory T cells and CD4 Naïve T cells.

Results reported in **Fig. 2e** and **Supplementary Fig. 1** support the inference that effector and cytotoxic immune cell types are less clearly differentiated as people age^11^, with stronger phenotypic convergence observed in males. Notably, the opposite phenomenon was true in Tregs and Naïve CD4 T cells, with both cell types undergoing clear gain of cell-type identity markers in elderly males and females (**Fig. 2f)**. FastDE additionally demonstrated that young (18-40 years old) and middle-aged (41-60 years old) adult cell-types closely resemble the ensemble profile, thus providing a reasonable approximation of the marker identities amongst these 19 predominant immune cell types in human blood. Complete lists of markers are provided in **Supplementary Tables 2**,**3**.

We replicated this result by applying the same FastDE workflow to an independent million-cell dataset, the OneK1K PBMC atlas of ∼1000 cells from each of 982 Australian adults^12^. Despite observing a slightly older age pyramid, the core findings that effector immune cell types have less divergent gene expression profiles while Tregs and Naïve CD4 T cells undergo an age-related gain in cell-type identity in the elderly population, over the age of 60, replicated (**Supplementary Fig. 2**).

Computational performance and scalability of FastDE was further demonstrated by contrasting run times for the complete OneK1K PBMC atlas and the compendium PBMC dataset. The DEG computation with Seurat required 6.5 hours on one CPU core and 1.25 hours on 64 cores for the OneK1K dataset with 1.2 million cells, while FastDE computed the same in 8 minutes on one core and 33 seconds on 64 cores. The compendium PBMC dataset includes 2.7 million cells and exceeded R’s two billion non-zero elements limit for sparse matrices, thus was not loadable in Seurat. FastDE was able to complete the computation in 14 minutes on one core and 45 seconds on 64 cores.

In summary, implementation of a series of optimization steps during the ranking of gene expression for marker detection, and recoding Seurat in C++ to utilize sparse matrices, facilitates significant speed-up of the workflow and analysis of million-cell datasets. FastDE software is freely available at GitHub. It is expected to support a wide range of applications from marker detection in cell states and types to quantitative comparison of differential expression in expansive single cell atlases.

## Acknowledgements

EC is supported by training grant 1T32EB021962-01A1 “Research Training Program in ImmunoEngineering” awarded to Prof. Julia Babensee at Georgia Tech. We are grateful to Joseph Powell for discussions relating to the OneK1K dataset, and to Prof. Srinivas Aluru for support of TP, and a GTRI-IRAD DE00019648 for support of TP, MA, and SC.

