## Supplementary material for "Acceleration of Large-Scale Single Cell Differential Gene Expression Analysis with FastDE": Methods and Supplemental Data

### FastDE

FastDE is implemented as an R package with C++ for compute-intensive components. The C++ components include Wilcoxon Rank Sum test, find markers, fold change, and supporting functions such as sparse matrix transpose and sums. The R FastDE package provides R language binding for the C++ functions as well as high level functions meant as direct replacement of the Seurat "FoldChange" and "FindMarkers" functions. We used the "cpp11" package (https://CRAN.R-project.org/package=cpp11) for C++ to R language binding for its simple semantics and light weight mapping compared to the "Rcpp" package (https://CRAN.R-project.org/package=Rcpp), particularly for read-only function parameters. Parallelization is accomplished using OpenMP in the C++ implementation. FastDE's core C++ implementation is available from https://github.com/tcpan/fastde-cpp. The FastDE R package utilizes the C++ implementation and is available as source from https://github.com/tcpan/fastde and has been submitted to the CRAN R-repository.

(1) Large Sparse Matrix Support

R's native sparse matrix class "dgCMatrix" uses compressed sparse column representation that internally consists of the value array, *X*, the index array, *I*, and the offset array, *P*. We adopt the organization that each row represents a cell, and each column represents a gene. The value array contains the non-zero values of the sparse matrix corresponding to the gene expression counts. The index array is the same size as *X* and contains the row indices of the elements, thus the cell ids. The offset array marks the positions of the first non-zero elements of each matrix column within the value and index arrays and has size equal to the number of genes plus one.

The offset array in dgCMatrix has element type integer, whose binary representation limits the number of non-zeros elements to $2^{32}$ or approximately two billion. We define "large sparse matrix" as those with more than $2^{32}$ nonzero elements. FastDE includes a new class, "dgCMatrix64", that supports a large sparse matrix by using a 64-bit floating point number offset array to support up to $2^{53}$ or over 8 quadrillion non-zero elements based on the IEEE 754 standard. All FastDE functions are defined to support both dgCMatrix and dgCMatrix64, and additionally the *X*, *P*, and *I* arrays directly, as input. FastDE further defines parallelized implementations of sparse matrix functions for matrix transposition, concatenation, row- and column-wise summation, and dense matrix conversion.

As R does not natively support loading large sparse matrices as dgCMatrix, FastDE supports two alternative approaches, either loading the *X*, *P*, and *I* arrays then creating a dgCMatrix64 instance, or loading R-compatible small sparse matrices as a list of dgCMatrix instances and combining them to form a dgCMatrix64 instance via FastDE's "cbind" and "rbind" matrix concatenation functions.

(2) Wilcoxon Rank Sum Test Algorithm and Implementation

The Wilcoxon Rank Sum test algorithm is briefly described in the main text. Our algorithm implementation is summarized in **Supplementary Fig. 3**. The algorithm accepts as input the *X*, *I*, and *P* arrays as described previously, as well as a *CellLabels* array containing the class labels of each cell and has length equal to the number of cells in the dataset (line 1).

The algorithm first counts the number of cells for each class label (lines 2-4). This is simply done via a linear traversal of *CellLabels* and accumulating the count in the *label_counts* hash table. The linear read of the *CellLabels* array makes optimal use of cache lines and memory access latency is reduced through CPU hardware prefetching, while the random memory writes are against a small hash table that is likely fully resident in cache. This step has complexity of $O(n)$ where *n* is the number of cells.

Next our algorithm computes for each gene the rank sums and p-values of every class label in a one-versus-all-others fashion (lines 7-15). The consecutive elements in the *P* array indicate the inclusive starting (***s***) and exclusive ending (***e***) position of elements belonging to a matrix column. For each column, we sort the elements (***sort***, line 10), compute the rank sum for each class as well as the tie correction (***sum_ranks***, line 11), and the U statistics and p-value of each class (***compute_pval***, lines 12-14). The algorithm returns a p-value matrix, where each row corresponds to a class label and each column a gene. A p-value in the matrix indicates whether the expression counts of the cells with a particular class label are statistically different than those of other cells for a particular gene. The differentially expressed genes for each class can then be identified based on the p-values using existing Seurat logic.

The ***sort*** operation (line 10) first creates a temporary array of expression values and cell class label pairs for the non-zero column elements. The class labels are extracted from the *CellLabels* array based on cell ids in the *I* array between positions ***s*** and ***e***. The traversals are linear and therefore cache efficient. The temporary array is then sorted by value followed by class label and returned as the *sorted* array. In this operation, given the expected data sparsity, random memory access during the sorting operation is likely to primarily involve data in cache. The numbers of zero-valued cells per class are also computed by subtracting the number of non-zero counts per class from the *label_counts* hash table and returned as the *label_zeroes* hash table. This operation is dominated by the sort complexity, which is expected to be approximately $O(|X| \; log(m))$ where |X| is the number non-zero matrix elements and *m* is the average number of nonzero elements in a column.

The *sorted* array is then traversed linearly to compute the per-class rank sums using the ***sum_ranks*** operation (line 11) according to the equation $R_1 = \sum_{i=1}^{n} I(i) r_i$, $I(i) = 1$ if $i$ in the class, 0 otherwise. We compute the in-class rank sums, $R_1$, of all classes explicitly, and the "all-except-class" rank sums from $R_1$ via a constant-time computation, $R_2 = \frac{N(N+1)}{2} - R_1$. Instead of explicitly storing the ranks as an array, they are implicitly tracked during the sorted array traversal. For each non-zero element in the sorted array, its rank in the sorted list is accumulated in the *rank_sums*[*label*] hash table element. We leverage the fact that all the implicit zero-elements would have occupied a contiguous block in the sorted array, and that they would share the same rank, to compute the contribution of the zero-elements in constant time. As the gene expression count matrix contains only positive values, we can assume that all zero-elements would have been placed at the beginning of the *sorted* array, and therefore the first rank must equal to the number of zero-elements, or alternatively cell counts minus the number of non-zero elements in the column.

Where there exist ties in the *sorted* array, the Wilcoxon Rank Sum Test uses the median rank of the ties, i.e. the average of the minimum and maximum positions of the ties. Scanning ahead to find the maximum position of a tie in order to compute the median rank, then iterating over the tied elements again to sum the ranks can introduce irregular, non-linear array access patterns, inactivates hardware prefetching, and increases memory access latency. Instead, we use a forward traversal pass to sum the minimum ranks including the ties, and a reverse traversal pass to sum the maximum ranks including the ties. Halving the sums in the *rank_sums* hash table produces the desired sum of median ranks. During the forward pass, we also compute *tie_sum*, to be used in the next step for tie correction, based on the expression $\sum_{i=1}^{k} t_i^3 - t_i$, where $t_i$ is the number of elements in a particular tie. We note that $t_i$ is a property of the tie and not dependent on the class labels, therefore the *tie_sum* for a column is common for all classes. The ***sum_ranks*** operation primarily involves linear traversal of the sorted array and therefore has complexity $O(|X|)$.

The *rank_sums* for all classes are then converted into the U statistics of the Wilcoxon Mann Whitney test in the ***compute_pval*** operation based on the equations $U_1 = R_1 - \frac{n_1(n_1+1)}{2}$ and $U_2 = n_1 n_2 - U_1$ for each class. The in-class cell counts $n_1$is retrieved from the *label_counts* hash table, while the out-of-class cell count is computed as $n_2 = n - n_1$, $n$ being the total number of cells. The U statistics are converted to z-scores with tie and continuity corrections. For two-tailed tests, $z = \frac{\max(U_1, U_2) - m_U - 0.5}{\sigma_U}$, where $m_U = \frac{n_1 n_2}{2}$ and $\sigma_U = \sqrt{\frac{n_1 n_2}{12}\left((n+1) - \frac{tie\_sum}{n(n-1)}\right)}$. From the z-scores, p-values are derived via the normal distribution CDF using the C function $erfc\left(\frac{z}{\sqrt{2}}\right)$ for a two-tailed test. The computation for each p-value requires constant time, and a p-value is computed for each class and each gene, therefore the time complexity for the operation is $O(cg)$, where *c* is the number of classes and *g* is the number of genes. Overall, the algorithm's complexity is dominated by that of the ***sort*** operation.

Our implementation is fully parallelized for multi-core, multi-thread environments. The matrix is partitioned into blocks of columns (line 7 in **Supplementary Fig. 3**) and the p-value submatrix is computed in blocks by each thread using OpenMP parallel sections.

The "FoldChange" operation is implemented similarly using a hash table to store intermediate sums for each class. For each gene, the class-versus-rest fold change values is then computed from the intermediate sums.

(3) <u>Seurat compatible FastFindAllMarkers function</u>

Seurat's "FindAllMarkers" implementation operates on a gene column for one class at a time, using a nested loop that iterates over the cell classes in the outer loop, and processes the gene columns in the inner loop using the "FoldChange" and "FindMarkers" functions. In this configuration, the sorted column therefore cannot be reused for multiple classes. FastDE's implementation of "FindMarkers" and "FoldChange" incorporates the entire nested loops and inverts the nesting order, iterating over the gene columns in the outer loop, and simultaneously computes the rank sums and p-values for all cell classes in the inner loop. FastDE's "FastFindAllMarkers" function invokes its accelerated FindMarkers and FoldChange functions when "fastwmw" is specified as the algorithm, and delegates to Seurat's "FindAllMarkers" function for Seurat-predefined algorithms.

(4) <u>Evaluation and validation</u>

The performance evaluation was conducted using a system with four Intel(R) Xeon(R) E7-8870 v3 CPUs each with 18-cores for a total of 72 CPU cores and 1TB of DDR4 ECC memory. The operating system used is Ubuntu 18.04.6 LTS. A 224GB NVME SSD was used to host the datasets and the pipeline outputs.

R version 4.2.2 and Seurat 4.3.0 were used in the Seurat and FastDE evaluation. R packages required for Seurat and FastDE were installed through the R Bioconductor and devtools packages. A full list of package versions is in **Supplementary Table 4**. All packages were compiled with compilation flags "-O2 -mtune=native -march=native -fopenmp", specified in the R user-level makevars file. Default OpenBLAS library provides low level linear algebra routines.

FastDE's FoldChange, FindMarkers, and Wilcoxon Rank Sum Test implementations were validated using randomly generated sparse matrices. The outputs of the Wilcoxon Rank Sum test are compared to the corresponding R functions, and the mean, standard deviation, and range of the value differences, as well as root mean square error were computed and shown to be within R's default tolerance of approximately $1.5\text{x}10^{-8}$. FastDE and Seurat's "FoldChange" functions show higher variability due to differences in R and C++'s rounding policy for negative numbers.

PBMC datasets were chosen with progressively higher cell counts for assessing the scalability of the computational performance of FastDE and Seurat. Seven datasets with approximately 3K, 6K, 8K, 10K, 33K, 68K, and 600K cells were included in the evaluation. **Supplementary Table 5** lists the provider and URL from which to download the datasets. In each case, filtered or normalized matrices were used.

The typical Seurat single cell analysis pipeline, similar to the Seurat PBMC 3K tutorial (https://satijalab.org/seurat/articles/pbmc3k_tutorial.html), was used for the evaluations. The pipeline steps include data set loading, dimensionality reduction, neighbor finding and clustering, visualization, and differential gene expression analysis. Default parameters were used for all the steps, including algorithm choices: PCA for dimensionality reduction, Louvain for clustering, and UMAP for visualization. Run times for the pipelines, excluding file I/O times, as well as the differential gene expression analysis step were captured using the tictoc package in R. Each run was repeated three times and the average reported in **Supplementary Table 1**. The table shows the scalability and performance of FastDE compared to Seurat.

### Code Availability

FastDE code is available at GitHub (https://github.com/tcpan/fastde-cpp, https://github.com/tcpan/fastde) and the R package has been submitted to CRAN.

### Human scRNA-seq analysis

Our discovery human immune atlas was composed of ~2.7 million PBMCs (peripheral blood monocyte immune cells) from 679 individuals sampled in 36 datasets collected from public repositories as listed in **Supplementary Table 6**, with data downloaded directly from the Gene Expression Omnibus (GEO; https://www.ncbi.nlm.nih.gov/geo), European Nucleotide Archive (ENA; https://www.ebi.ac.uk/ena), or Single Cell Portal (SCP; https://singlecell.broadinstitute.org/single_cell). The selected studies were published between April 2018 and November 2022 and the incorporated datasets were mostly generated on the commonly employed 10x Chromium platform[13] (10x Genomics, 34 studies), although one study used Seq-well technology[14] (Gierahn et al., 2017). In our selection process, we prioritized studies that employed similar protocols for processing samples and generating data, such as sequencing of PBMCs. From studies contrasting healthy versus disease, we exclusively included the healthy control samples of the published data. We did not exclude studies that applied flow cytometry-based cell-sorting prior to sequencing. The integrated discovery PBMC atlas was generated and analyzed using the following steps: (1) pre-processing and filtering individual scRNA-seq datasets separately from healthy human blood; (2) integrating and clustering PBMCs from each dataset to generate a single healthy human PBMC atlas; (3) annotation of cell clusters; and (4) differential expression analysis. These four steps are described in detail in the following sections.

(1) Pre-processing and filtering individual scRNA-seq datasets

Single cell transcriptomics datasets, enriched in PBMCs and available as processed CellRanger files[13], were collected from public repositories as listed in **Supplementary Table 6**. A total of 35 scRNA-seq datasets representing 679 healthy subjects from multiple ancestries were analyzed individually incorporating metadata listed in **Supplementary Table 7**. To ensure comparability for every individual dataset, we only retained genes found in the Ensembl human (GRCh38) gene model, and implemented the same basic Seurat single-cell analysis pipeline (version 4.1.1[15]) in R (version 4.2.1). Specifically, for each dataset low quality cells with a high percentage of mitochondrial gene counts (>~5–20%, depending on proportion of outlier removal for each dataset) and with <500 measured genes were excluded. To mitigate potential doublet inclusion, cells with UMI count above 40,000 and detected genes above 5,000 were removed. This led to a working dataset of 2,742,544 single cells. In addition, the OneK1K atlas[12] was imported without modification as a validation set, for which metadata is provided in **Supplementary Table 8**.

(2) <u>Dataset integration for the healthy human PBMC discovery atlas.</u>

Next, the individual Seurat objects were processed to identify the 19 major cell clusters identified in the Azimuth routine of Seurat v4[15], and then merged one-by-one into a single healthy-state human PBMC object. After filtering, data in each cell was log normalized using Seurat's 'NormalizeData' function (method = 'LogNormalize', scale.factor = 10,000), the 2,000 most variable genes were identified, and the 'ScaleData' function was used to scale and center the gene expression matrix after regressing out the heterogeneity associated with cell cycle and mitochondrial contamination. For each dataset, the number of principal components used for neighborhood graph construction and dimensional reduction was set at 20. Cell clusters marked by the canonical marker genes for platelets (PPBP) and erythrocytes (HBB) were discarded. All individual datasets devoid of platelets and erythrocytes were then used for integration to create our core discovery PBMC atlas comprising data from 35 studies. Batch effect correction was performed on these processed, merged objects using Seurat's reciprocal PCA ('RPCA')[16] with study ID as the batch term with all other parameters as per default. After integration, Uniform Manifold Approximation and Projection (UMAP)[17] visualization indicated cells from different studies were well mixed into the shared space (**Fig. 2d;** cf **Supplementary Fig. 4** for OneK1K validation).

(3) <u>Annotation of cell clusters</u>

To simplify the analysis, prior annotations were used as a reference to annotate each cell in each dataset. If prior annotations were not available, cellular identity was determined by finding DE genes for each cluster by the 'FastFindAllMarkers' function of the FastDE package (test.use= 'wilcox', min.pct=0.1, logfc.threshold=.5). We applied two complementary methods to confirm the cluster annotations shown in Fig 1C. We first compared the top ranking differentially expressed genes of query clusters to the well-characterized cell type-specific genes from previous datasets listed in **Supplementary Table 9**. We then assessed cluster annotation by applying the R package Azimuth[15] which compares the transcriptome of each single cell to reference PBMC transcriptomic datasets to validate cellular identity. Feature plots in **Supplementary Fig. 5** illustrate some key clusters. For the OneK1K dataset[12] we accepted the Azimuth annotations of predicted cell type identities as published on the CZI (http://cellxgene.cziscience.com) website from which it was downloaded.

(4) Fast differential expression (DE) analysis for clusters and groups

A crucial decision with respect to DE analysis is whether to preserve the intrinsic makeup of cells and samples within the dataset or to use a more structured design in which the numbers of cells and individuals are downsampled and balanced prior to performing the contrast of interest. We chose the latter approach based on pilot analyses reported in **Supplementary Fig. 6** exploring the impacts of imbalanced sex ratio, sample number, and cell number. The results show the total number of sex-specific DEGs (in memory B cells) varies according to the number of cells (test 3); and the ratio of male to female samples (test 1), but not the total number of cells (test 2). In our analyses, differences in the number of cells (test 3: 5K, 12K, or 25K) showed substantial effect on the total number of sex-specific DEGs, with smaller number of cells tending to show higher DEG numbers. On the same scale, the balance of samples (test 1: 25 females:75 males or 50 females:50 males) showed a similar magnitude of effect on DEGs detected, with nearly a 2-fold increase in the total number of DEGs when samples were unbalanced as compared to balanced even after adjusting for cell number. Analysis of the total number of samples (test 2: 100 or 200 males and females) exhibited little change in the total number of DEGs, indicating that there is no tendency for genes that are differentially expressed to be subject to sample size differences.

Thus, to account for any potential effect of variable group-specific cell numbers on gene expression, cells were randomly downsampled across age and sex subgroups so that the total cell-type count was the same across each sex decile. From the parent Seurat objects of both the

Discovery and the OneK1K atlases, healthy subjects were divided into twenty groups (10 age deciles for males and 10 age deciles for females) according to sex and the age distributions for each cohort (**Supplementary Fig. 7**). Note that the differences in decile ranges mostly reflect the differences in the age pyramids of the Discovery and OneK1K cohorts, shown in **Supplementary Fig. 8**.

Differential expression analysis for each cell type versus all other cells within a sex decile was performed using the Wilcoxon-test as implemented in the ‘‘FastFindAllMarkers’’ function of the FastDE package. DEGs were identified using the following criteria: (1) a logfold change >0.50, (2) adjusted p-value <0.05, (3) transcripts detected in >10% of cells in either test group. Plots for the differentially expressed genes were generated by customized R code using ggplot2 (v3.3.3, R package).

### SUPPLEMENTARY MATERIAL

### SUPPLEMENTARY FIGURES

**Suppl. Fig. 1.** Extended Association of Cell Type Identity with Age in the Discovery Atlas.

**Suppl. Fig. 2.** Replication Cell Type Identity Associations with Age in the OneK1K Atlas.

**Suppl. Fig. 3.** Wilcoxon Rank Sum Test Algorithm.

**Suppl. Fig. 4.** UMAP visualization of OneK1K Atlas.

**Suppl. Fig. 5.** Feature Plots of Canonical Markers of Immune cells in the Discovery Atlas.

**Suppl. Fig. 6.** Justification for dividing samples by decile rather than age group.

**Suppl. Fig. 7.** Age distribution for deciles of the PBMC Data Atlases.

**Suppl. Fig. 8.** Age pyramids for donors to the PBMC Data Atlases.

### SUPPLEMENTARY TABLES

**Suppl. Table 1.** Summary statistics for workflow acceleration using FastDE.

**Suppl. Table 2.** Complete list of cell-type specific markers identified by FastFindAllMarkers.

**Suppl. Table 3.** Cell-type specific markers identified by FastFindAllMarkers for each age and sex subgroup.

**Suppl. Table 4.** R package versions used for evaluation of computational performance.

**Suppl. Table 5.** Datasets used for evaluation of FastDE computational performance.

**Suppl. Table 6.** Characteristics of studies included in the discovery atlas.

**Suppl. Table 7.** Metadata relating to healthy subjects in the discovery atlas.

**Suppl. Table 8.** Metadata relating to healthy subjects in the OneK1K validation atlas.

**Suppl. Table 9.** Classical markers used for cell type identification.

**Supplementary Table 1. Summary Statistics for Workflow Acceleration using FastDE**

#### A. Single core Processing

| Dataset (k cells) | FastDE | | Seurat v4.3 | | Speed up | |
|---|---|---|---|---|---|---|
| | FindMarkers (sec) | Pipeline (sec) | FindMarkers (sec) | Pipeline (sec) | FindMarkers | Pipeline |
| 3 | 1.56 | 24.80 | 88.37 | 111.46 | 56.57 | 4.49 |
| 6 | 2.08 | 42.82 | 159.04 | 199.86 | 76.41 | 4.67 |
| 8 | 4.90 | 57.67 | 477.02 | 529.69 | 97.29 | 9.19 |
| 10 | 9.42 | 60.15 | 1839.36 | 1890.08 | 195.28 | 31.42 |
| 33 | 9.31 | 177.90 | 1591.56 | 1760.17 | 170.95 | 9.89 |
| 68 | 13.30 | 388.77 | 2608.77 | 2985.35 | 196.20 | 7.68 |
| 600 | 233.58 | 2574.91 | 145565.11 | 148029.67 | 623.19 | 57.49 |

#### B. 64 Core Processing

| Dataset (k cells) | FastDE | | Seurat v4.3 | | Speed up | |
|---|---|---|---|---|---|---|
| | FindMarkers (sec) | Pipeline (sec) | FindMarkers (sec) | Pipeline (sec) | FindMarkers | Pipeline |
| 3 | 1.16 | 10.73 | 19.42 | 28.99 | 16.68 | 2.70 |
| 6 | 1.20 | 18.52 | 30.11 | 47.43 | 25.13 | 2.56 |
| 8 | 1.98 | 27.18 | 79.36 | 104.56 | 40.04 | 3.85 |
| 10 | 3.11 | 28.05 | 225.56 | 250.49 | 72.46 | 8.93 |
| 33 | 2.82 | 84.03 | 191.24 | 272.45 | 67.87 | 3.24 |
| 68 | 3.50 | 191.02 | 335.51 | 523.03 | 95.79 | 2.74 |
| 600 | 28.73 | 2389.61 | 2805.33 | 5166.20 | 97.63 | 2.16 |

The run times are reported in seconds and speed ups are calculated as ratios of Seurat's times to FastDE's times. The pipelines were executive using one core of a Xeon E7-8870 CPU for the FindMarkers function and the complete pipeline. Pipeline run times exclude file input and output times.

**Supplementary Table 4.** List of installed R packages and versions used during evaluation of computational performance.

| R package | version | R package | version | R package | version |
|---|---|---|---|---|---|
| nlme | 3.1-162 | matrixStats | 0.63.0 | spatstat.sparse | 3.0-1 |
| RcppAnnoy | 0.0.20 | RColorBrewer | 1.1-3 | httr | 1.4.4 |
| sctransform | 0.3.5 | tools | 4.2.2 | utf8 | 1.2.2 |
| R6 | 2.5.1 | irlba | 2.3.5.1 | KernSmooth | 2.23-20 |
| uwot | 0.1.14 | lazyeval | 0.2.2 | colorspace | 2.0-3 |
| sp | 1.6-0 | tidyselect | 1.2.0 | gridExtra | 2.3 |
| compiler | 4.2.2 | progressr | 0.13.0 | cli | 3.6.1 |
| spatstat.explore | 3.1-0 | plotly | 4.10.1 | scales | 1.2.1 |
| lmtest | 0.9-40 | spatstat.data | 3.0-1 | ggridges | 0.5.4 |
| pbapply | 1.7-0 | goftest | 1.2-3 | stringr | 1.5.0 |
| digest | 0.6.30 | spatstat.utils | 3.0-2 | pkgconfig | 2.0.3 |
| htmltools | 0.5.4 | parallelly | 1.35.0 | fastmap | 1.1.0 |
| htmlwidgets | 1.6.2 | rlang | 1.1.0 | shiny | 1.7.4 |
| generics | 0.1.3 | zoo | 1.8-11 | jsonlite | 1.8.4 |
| spatstat.random | 3.1-4 | ica | 1.0-3 | dplyr | 1.1.1 |
| magrittr | 2.0.3 | patchwork | 1.1.2 | Matrix | 1.5-3 |
| Rcpp | 1.0.10 | munsell | 0.5.0 | fansi | 1.0.3 |
| abind | 1.4-5 | reticulate | 1.28 | lifecycle | 1.0.3 |
| stringi | 1.7.12 | MASS | 7.3-58.1 | Rtsne | 0.16 |
| plyr | 1.8.8 | grid | 4.2.2 | parallel | 4.2.2 |
| listenv | 0.9.0 | promises | 1.2.0.1 | ggrepel | 0.9.3 |
| deldir | 1.0-6 | miniUI | 0.1.1.1 | lattice | 0.20-45 |
| cowplot | 1.1.1 | splines | 4.2.2 | tensor | 1.5 |
| pillar | 1.8.1 | igraph | 1.4.1 | spatstat.geom | 3.1-0 |
| future.apply | 1.10.0 | reshape2 | 1.4.4 | codetools | 0.2-18 |
| leiden | 0.4.3 | glue | 1.6.2 | data.table | 1.14.8 |
| png | 0.1-8 | vctrs | 0.6.1 | httpuv | 1.6.9 |
| polyclip | 1.10-4 | gtable | 0.3.1 | RANN | 2.6.1 |
| purrr | 1.0.1 | tidyr | 1.3.0 | scattermore | 0.8 |
| future | 1.32.0 | ggplot2 | 3.4.1 | mime | 0.12 |
| xtable | 1.8-4 | later | 1.3.0 | survival | 3.5-0 |
| viridisLite | 0.4.1 | tibble | 3.2.1 | cluster | 2.1.4 |
| globals | 0.16.2 | fitdistrplus | 1.1-8 | ellipsis | 0.3.2 |
| ROCR | 1.0-11 | SeuratObject | 4.1.3 | Seurat | 4.3.0 |

**Supplementary Table 5.** Datasets used for evaluation of FastDE computational performance

| Dataset (k cells) | Source | Source | URL |
|---|---|---|---|
| 3 | Filtered | 10X Genomics | https://www.10xgenomics.com/resources/datasets/3-k-pbm-cs-from-a-healthy-donor-1-standard-1-1-0 |
| 6 | Filtered | 10X Genomics | https://www.10xgenomics.com/resources/datasets/6-k-pbm-cs-from-a-healthy-donor-1-standard-1-1-0 |
| 8 | Filtered | 10X Genomics | https://www.10xgenomics.com/resources/datasets/8-k-pbm-cs-from-a-healthy-donor-2-standard-2-1-0 |
| 10 | Filtered | 10X Genomics | https://www.10xgenomics.com/resources/datasets/10-k-pbm-cs-from-a-healthy-donor-v-3-chemistry-3-standard-3-0-0 |
| 33 | Filtered | 10X Genomics | https://www.10xgenomics.com/resources/datasets/33-k-pbm-cs-from-a-healthy-donor-1-standard-1-1-0 |
| 68 | Filtered | 10X Genomics | https://www.10xgenomics.com/resources/datasets/fresh-68-k-pbm-cs-donor-a-1-standard-1-1-0 |

Each dataset consists of three files: the mtx matrix file, a barcodes.tsv file, and a genes.tsv file. The filtered or normalized version were selected.

**Supplementary Table 9 Classical Markers Used for Cell Type Identification**

| Cell type | Markers |
|---|---|
| B intermediate | MS4A1, TNFRSF13B, IGHM, IGHD, AIM2, CD79A, LINC01857, RALGPS2, BANK1, CD79B |
| B memory | MS4A1, COCH, AIM2, BANK1, SSPN, CD79A, TEX9, RALGPS2, TNFRSF13C, LINC01781 |
| B naive | IGHM, IGHD, CD79A, IL4R, MS4A1, CXCR4, BTG1, TCL1A, CD79B, YBX3 |
| Plasmablast | IGHA2, MZB1, TNFRSF17, DERL3, TXNDC5, TNFRSF13B, POU2AF1, CPNE5, HRASLS2, NT5DC2 |
| CD4 CTL | GZMH, CD4, FGFBP2, ITGB1, GZMA, CST7, GNLY, B2M, IL32, NKG7 |
| CD4 Naive | TCF7, CD4, CCR7, IL7R, FHIT, LEF1, MAL, NOSIP, LDHB, PIK3IP1 |
| CD4 TCM | IL7R, TMSB10, CD4, ITGB1, LTB, TRAC, AQP3, LDHB, IL32, MAL |
| CD4 TEM | IL7R, CCL5, FYB1, GZMK, IL32, GZMA, KLRB1, TRAC, LTB, AQP3 |
| Treg | RTKN2, FOXP3, AC133644.2, CD4, IL2RA, TIGIT, CTLA4, FCRL3, LAIR2, IKZF2 |
| CD8 Naive | CD8B, S100B, CCR7, RGS10, NOSIP, LINC02446, LEF1, CRTAM, CD8A, OXNAD1 |
| CD8 TCM | CD8B, ANXA1, CD8A, KRT1, LINC02446, YBX3, IL7R, TRAC, NELL2, LDHB |
| CD8 TEM | CCL5, GZMH, CD8A, TRAC, KLRD1, NKG7, GZMK, CST7, CD8B, TRGC2 |
| cDC | FCER1A, CST3, SERPINF1, HLA-DQA1, CLEC10A, CD1C, ENHO, PLD4, GSN, SLC38A1, NDRG2, AFF3 |
| pDC | ITM2C, PLD4, SERPINF1, LILRA4, IL3RA, TPM2, MZB1, SPIB, IRF4, SMPD3 |
| CD14 Mono | S100A9, CTSS, S100A8, LYZ, VCAN, S100A12, IL1B, CD14, G0S2, FCN1 |
| CD16 Mono | CDKN1C, FCGR3A, PTPRC, LST1, IER5, MS4A7, RHOC, IFITM3, AIF1, HES4 |
| NK | GNLY, TYROBP, NKG7, FCER1G, GZMB, TRDC, PRF1, FGFBP2, SPON2, KLRF1 |
| NK_CD56bright | XCL2, FCER1G, SPINK2, TRDC, KLRC1, XCL1, SPTSSB, PPP1R9A, NCAM1, TNFRSF11A |
| MAIT | KLRB1, NKG7, GZMK, IL7R, SLC4A10, GZMA, CXCR6, PRSS35, RBM24, NCR3 |

**Supplementary Fig. 1. Extended association of cell type identity with age in the discovery atlas**

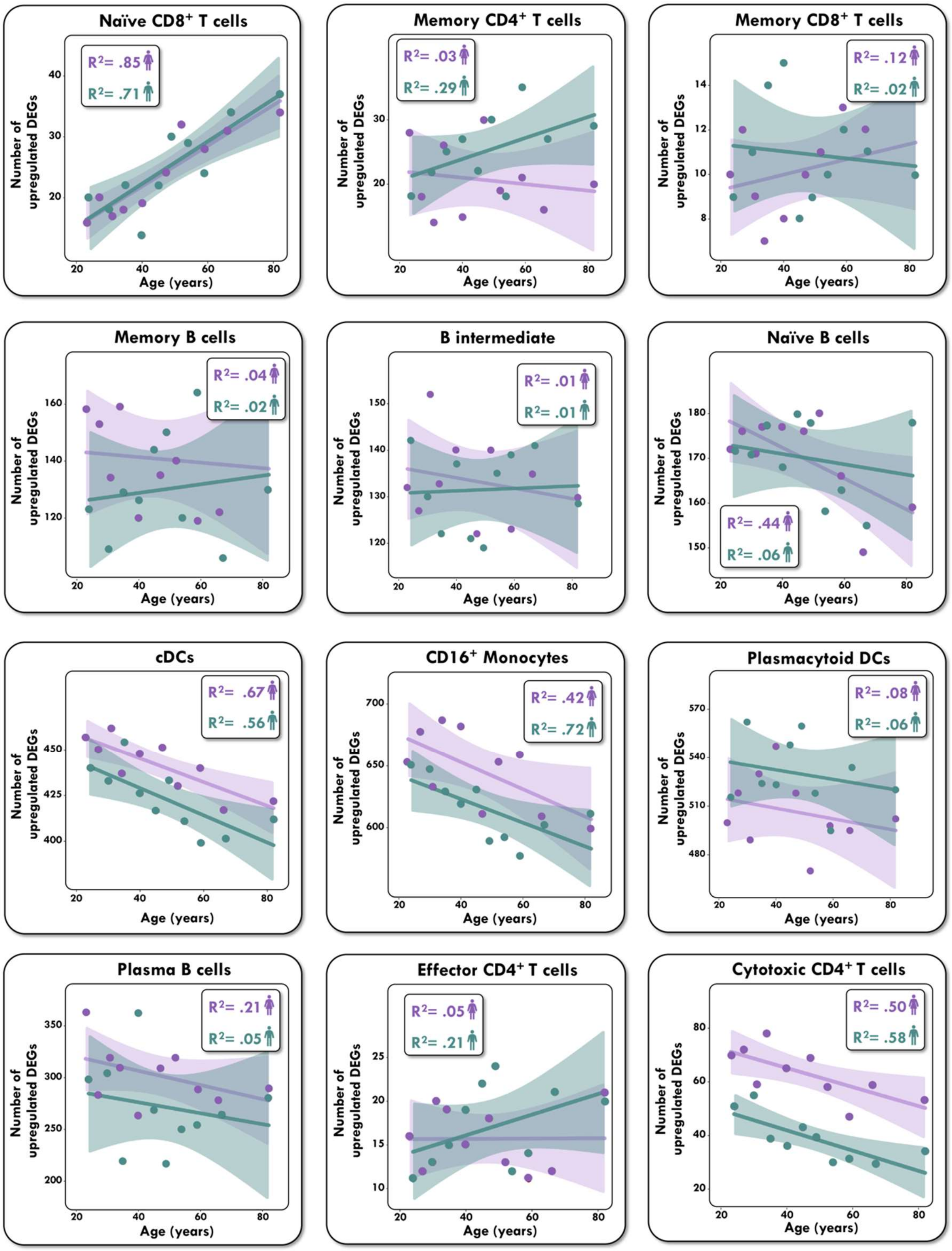

**Supplementary Fig. 2. Replication cell type identity associations with age in the OneK1K atlas**

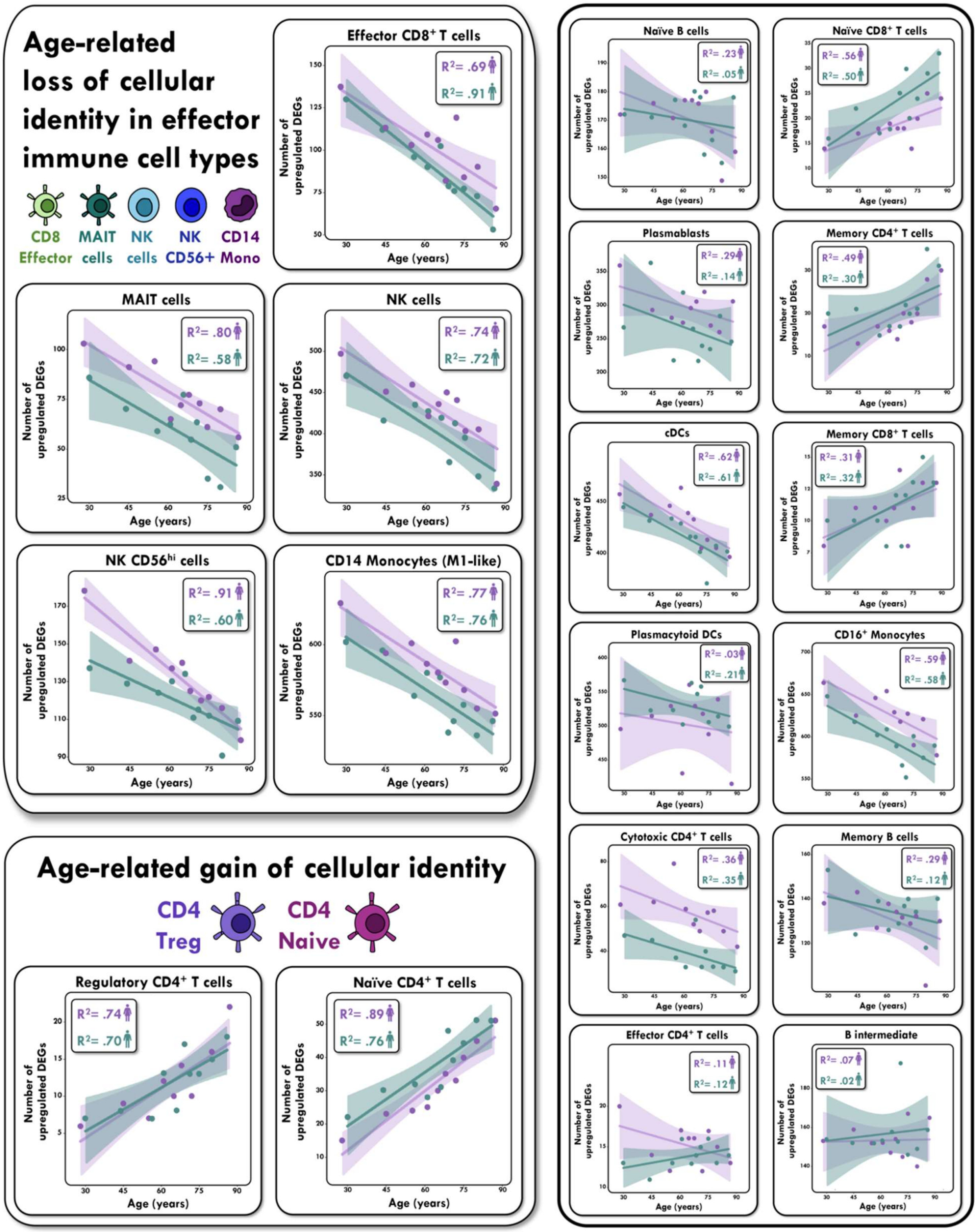

**Supplementary Fig. 3. FastDE Wilcoxon Rank Sum Index Algorithm**.

---

**Algorithm 1** Wilcoxon Rank Sum Test

---

**procedure** FAST_WILCOXON($X$, $I$, $P$, $CellLabels$)
  $label\_counts \Leftarrow$ hashtable
  **for** each $label$ in $CellLabels$ **do**
    increment $label\_counts[label]$
  **end for**
  $pval \Leftarrow$ zero matrix of dim $[|label_counts|, (|P| - 1)]$
  **for** $g \in \{1 \ldots (|P| - 1)\}$ **do**
    $s \Leftarrow P[g]$ ▷ column start
    $e \Leftarrow P[g + 1]$ ▷ column end
    $sorted$, $label\_zeroes \Leftarrow$ **sort**($X[s{:}e]$, $CellLabels[I[s{:}e]]$, $label\_counts$)
    $rank\_sums$, $tie\_sum \Leftarrow$ **sum_ranks**($sorted$, $label\_zeroes$)
    **for** each ($label$, $count$) $\in label_counts$ **do**
      $pval[label, g] \Leftarrow$ **compute_pval**($rank\_sums[label]$, $count$, $tie\_sum$
    **end for**
  **end for**
  **return** $pval$
**end procedure**

---

**Supplementary Fig. 4. UMAP visualization of OneK1K Atlas (Yazar et al 2022**).

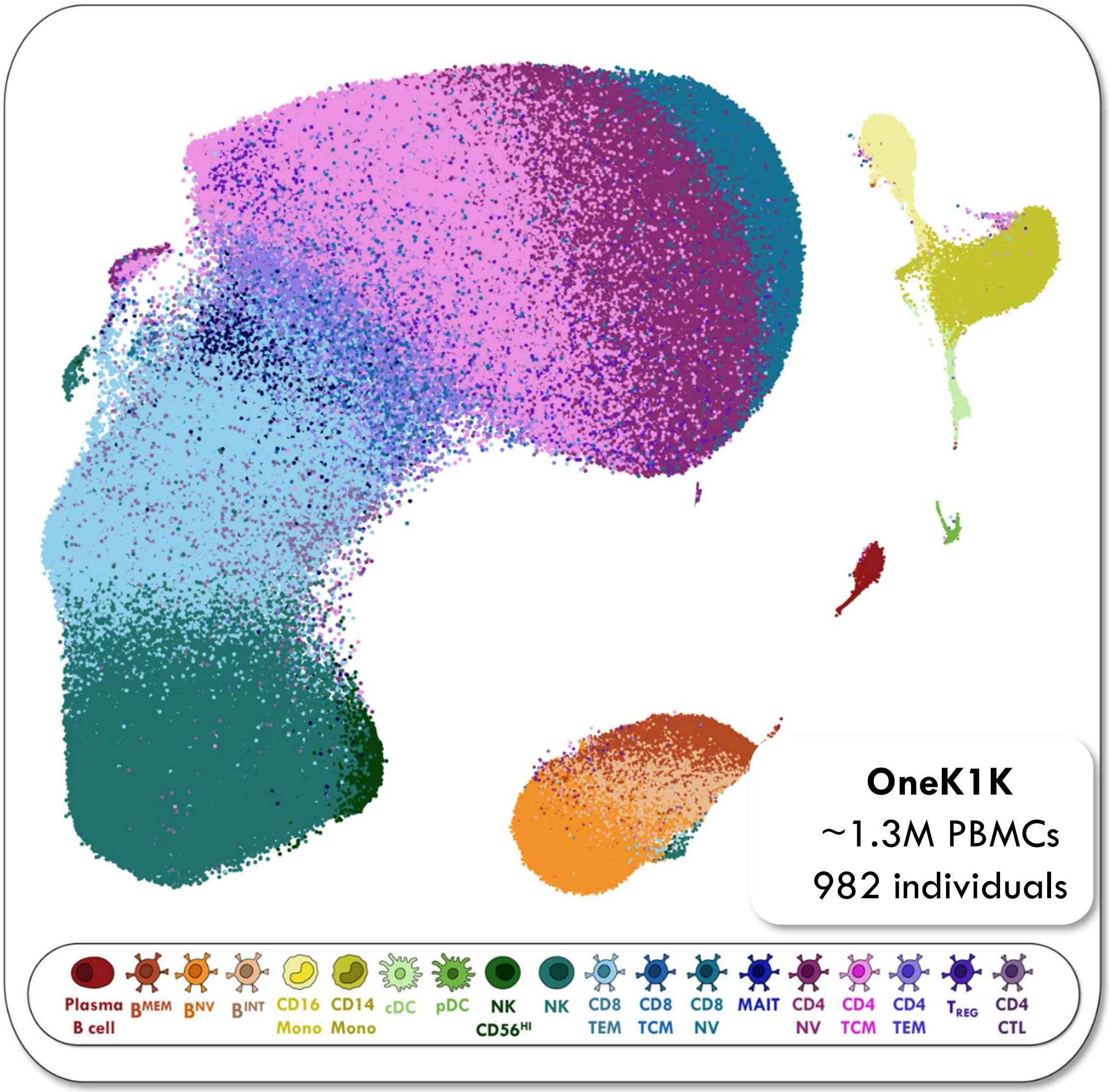

**Supplementary Fig. 5. Feature plots of canonical markers of immune cells in discovery atlas**

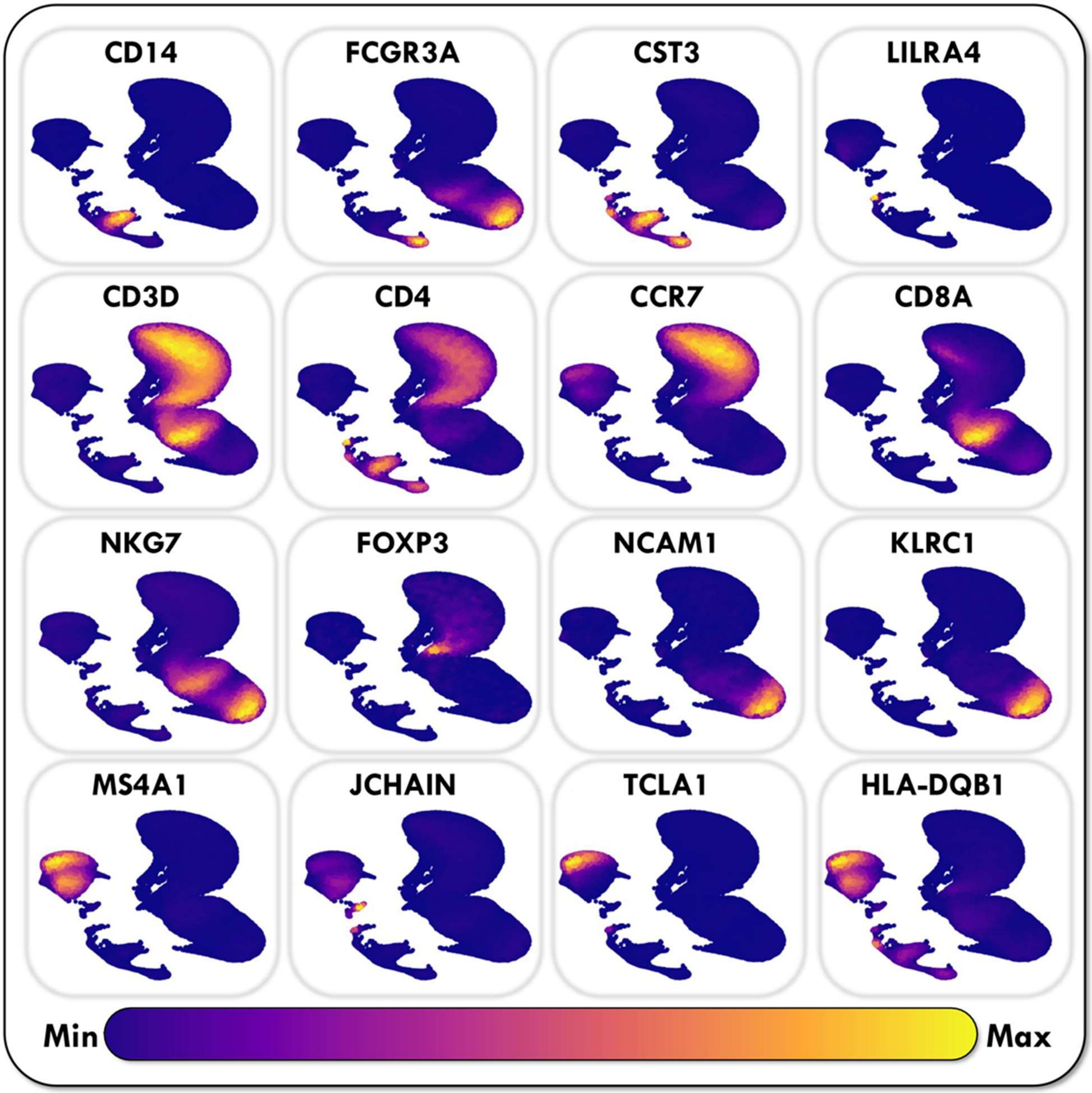

**Supplementary Fig. 6. Justification for dividing samples by decile rather than age group**

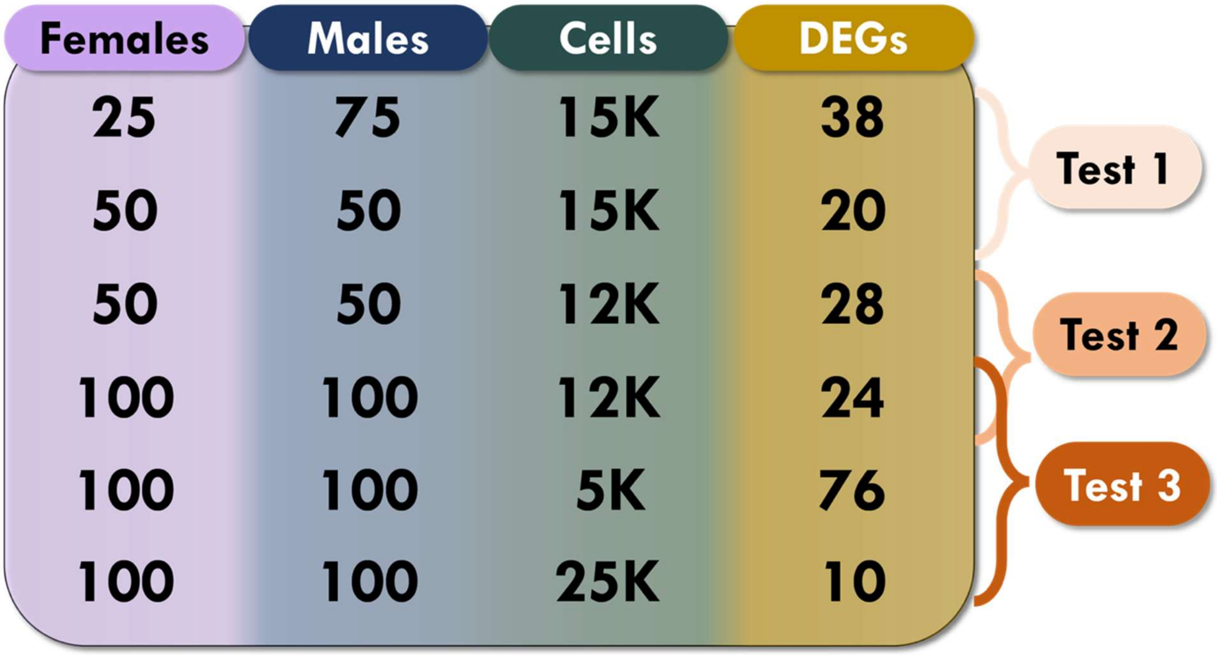


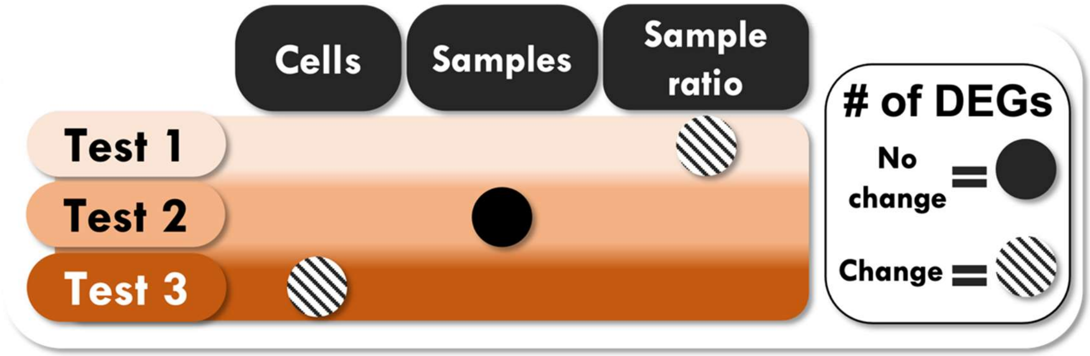

**Supplementary Fig. 7. Age distribution for deciles of the PBMC Data Atlases**

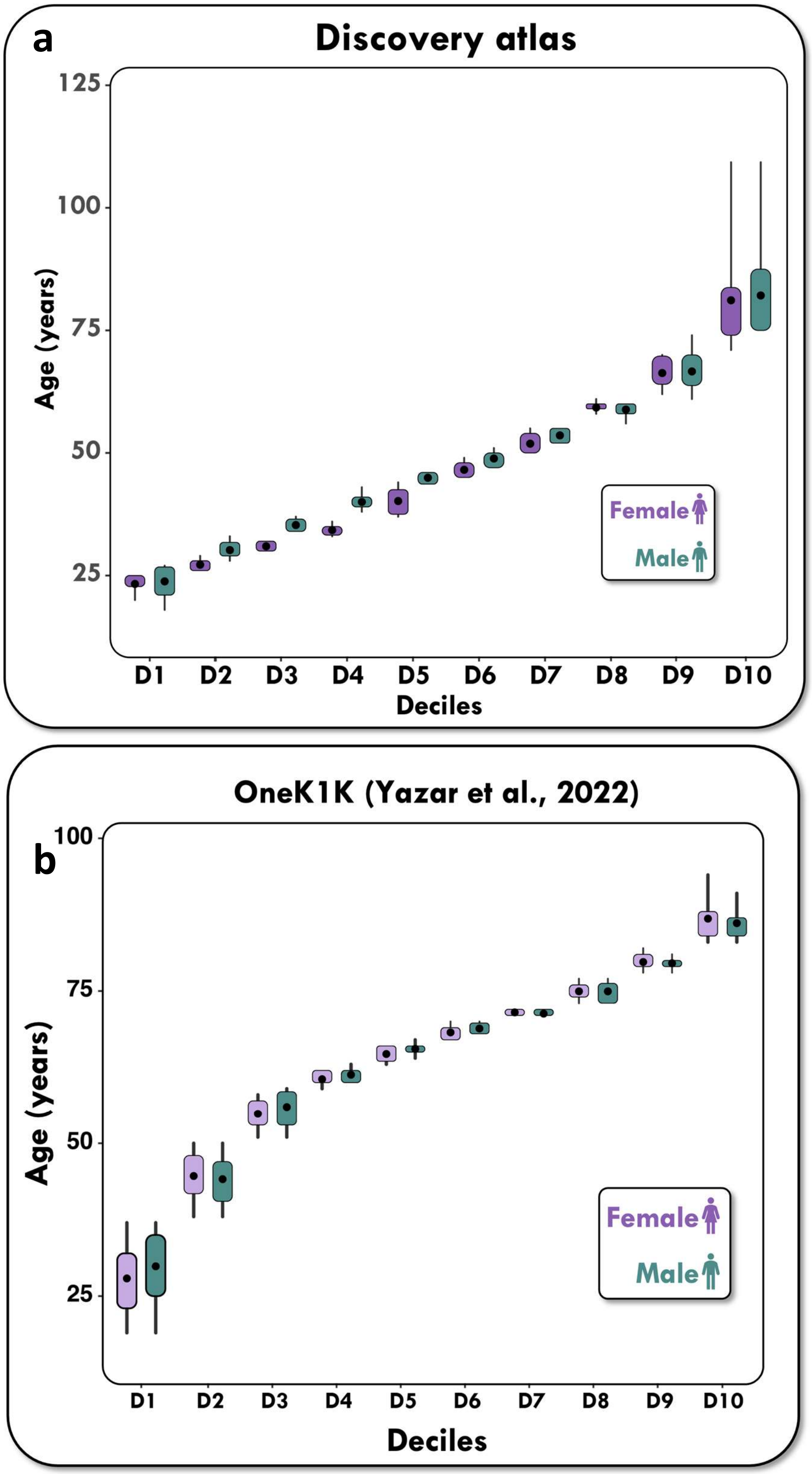

**Supplementary Fig. 8. Age pyramids for donors to the PBMC Data Atlases**

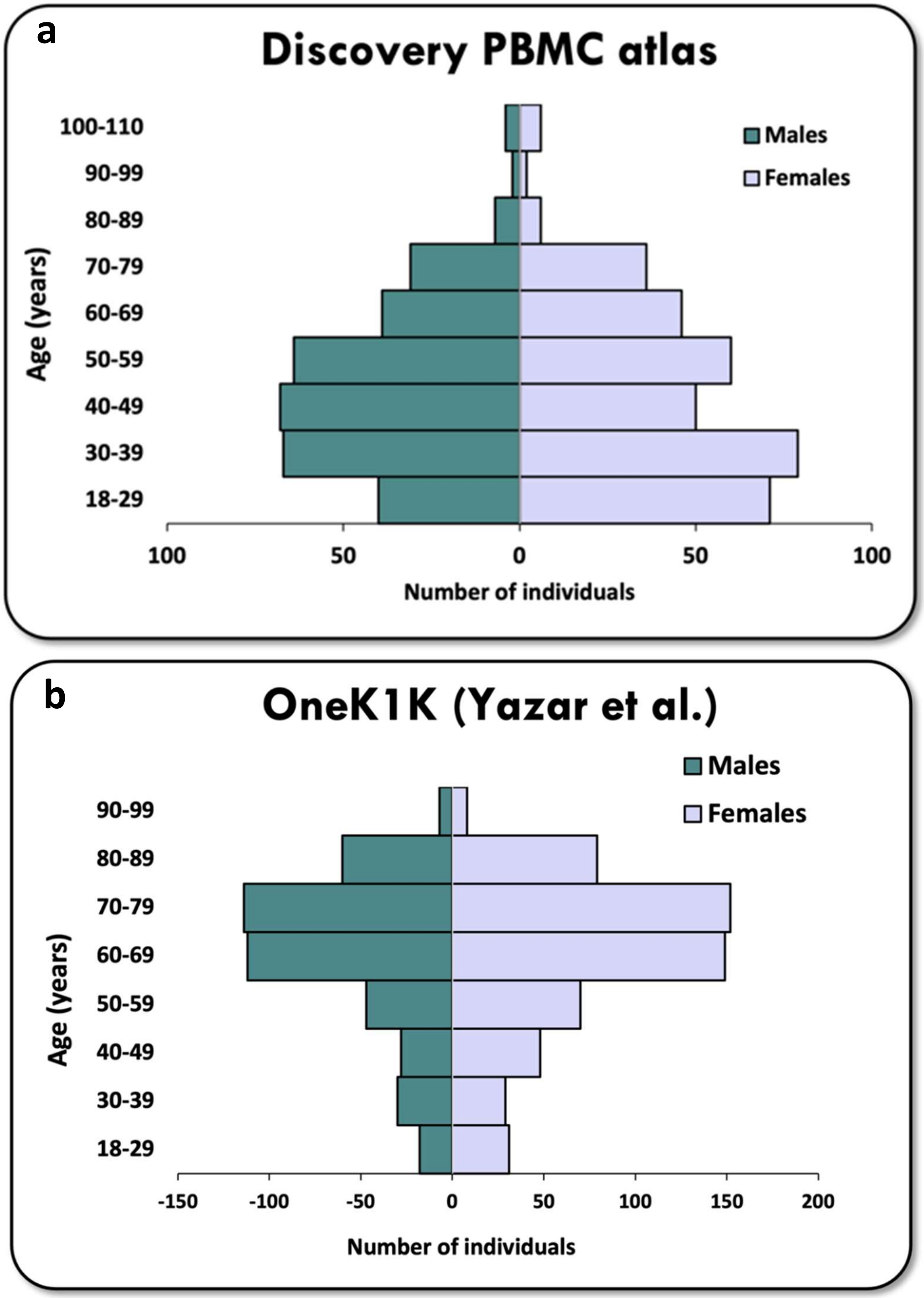
